# Using force data to self-pace an instrumented treadmill and measure self-selected walking speed

**DOI:** 10.1101/2019.12.18.870592

**Authors:** Seungmoon Song, HoJung Choi, Steven H. Collins

## Abstract

**Background:** Self-selected speed is an important functional index of walking. A self-pacing controller that reliably matches walking speed without additional hardware can be useful for measuring self-selected speed in a treadmill-based laboratory.

**Methods:** We adapted a previously proposed self-pacing controller for force-instrumented treadmills and validated its use for measuring self-selected speeds. We first evaluated the controller’s estimation of subject speed and position from the force-plates by comparing it to those from motion capture data. We then compared five tests of self-selected speed. Ten healthy adults completed a standard 10-meter walk test, a 150-meter walk test, a commonly used manual treadmill speed selection test, a two-minute self-paced treadmill test, and a 150-meter self-paced treadmill test. In each case, subjects were instructed to walk at or select their comfortable speed. We also assessed the time taken for a trial and a survey on comfort and ease of choosing a speed in all the tests.

**Results:** The self-pacing algorithm estimated subject speed and position accurately, with root mean square differences compared to motion capture of 0.023 m s^−1^ and 0.014 m, respectively. Self-selected speeds from both self-paced treadmill tests correlated well with those from the 10-meter walk test (*R* > 0.93, *p* < 1 × 10^−13^). Subjects walked slower on average in the self-paced treadmill tests (1.23 ± 0.27 m s^−1^) than in the 10-meter walk test (1.32 ± 0.18 m s^−1^) but the speed differences within subjects were consistent. These correlations and walking speeds are comparable to those from the manual treadmill speed selection test (*R* = 0.89, *p* = 3 × 10^−11^; 1.18 ± 0.24 m s^−1^). Comfort and ease of speed selection were similar in the self-paced tests and the manual speed selection test, but the self-paced tests required only about a third of the time to complete. Our results demonstrate that these self-paced treadmill tests can be a strong alternative to the commonly used manual treadmill speed selection test.

**Conclusions:** The self-paced force-instrumented treadmill well adapts to subject walking speed and reliably measures self-selected walking speeds. We provide the self-pacing software to facilitate use by gait researchers and clinicians.

## Background

Self-selected walking speed is one of the main performance indices of walking. It is the speed at which people normally choose to walk and is also known as preferred speed or comfortable speed. Walking speed determines the time required in achieving the primary goal of walking: getting to a destination. Healthy adults normally choose to walk at about 1.3 m s^−1^ although they can walk much faster (> 2.0 m s^−1^) [1]. Normal walking speed likely results from balancing many factors, including energy use, time spent in transit, appearance, and comfort. It has often been observed that self-selected walking speed is close to the speed that minimizes metabolic energy consumption [2, 3] or muscle fatigue [4] in traveling a unit distance. Self-selected walking speed also has been emphasized as a promising measure to assess physical health. For example, walking speed is a good predictor of health status and survival rate in older adults [5, 6] and a useful measure for rehabilitation progress [7].

There are different ways to measure self-selected walking speeds. A standard method commonly used in physical therapy and gait studies is the so-called 10-meter walk test [8, 9]. In a 10-meter walk test, subjects are instructed to walk at their comfortable speed across a 15∼20 m walkway, and the time taken to traverse the middle 10 m section is measured with a stopwatch to calculate self-selected walking speed. This process is often conducted multiple times then averaged for reliable measurements. Another common way of measuring self-selected speed is by asking subjects to manually select their comfortable speeds while walking on a treadmill that changes from slow to fast or fast to slow speeds [10, 11, 12]. Measuring comfortable speeds on a treadmill is useful for certain cases, such as collecting data in a treadmill-based gait laboratory [13] and studying assistive technologies with immobile systems [14]. On the other hand, this manual selection process requires the subjects to walk at various speeds, which can be time consuming, and to consciously distinguish comfortable from uncomfortable treadmill speeds, which can be confusing for those who are not familiar with walking on a treadmill.

Self-paced treadmills can also be useful in measuring walking speed. A treadmill that can seamlessly adapt to a subject’s walking speed can provide an overground-like walking environment and can compensate for shortcomings in the manual speed selection approach. Self-pacing controllers typically consist of two parts, usually treated independently. The first estimates the subject’s speed and position. The second controls treadmill speed based on the estimation. The treadmill speed is typically controlled to match subject speed and to keep the subject in the middle of the treadmill [15, 16]. Various approaches of estimating subject speed and position have been used. One approach is to use a marker-based optical motion capture system [17, 16, 18], which is widely used in research laboratories as a part of a commercial virtual reality package [19]. Researchers have evaluated these motion capture based self-paced treadmills by comparing kinematic and kinetic gait features collected on the self-paced treadmill to those during fixed speed treadmill walking [16] and overground walking [18]. In addition, these self-paced treadmills have been used in rehabilitation research for children with cerebral palsy [20, 21], individuals with chronic stroke [22], and individuals with transtibial amputation [23]. Other approaches with low-cost sensors or simpler hardware have been proposed as well, such as using a marker-free infrared-based motion sensor [24], an ultrasonic distance sensor [25], a harness with force sensors [26], and force plates on an instrumented treadmill [15].

A self-pacing controller using force-plate data from an instrumented treadmill is attractive because it does not require additional hardware or instrumentation. Feasel and colleagues [15] have proposed such a controller and used it to separately control the belts on a split-belt treadmill for asymmetric gait. They calculated the ground reaction forces and center of pressure from the force-plate data and combined them with a Kalman filter to track walking speed. The study focused on testing the feasibility of improving gait symmetry in hemiparetic patients with a virtual environment that integrated the self-paced treadmill and a visual scene. Although they reported that the hemiparetic patients self-selected to walk at speeds comparable to their overground speeds, a more thorough evaluation of self-selected walking speed on this type of self-paced treadmill would improve our understanding of its efficacy.

Various aspects of a walking speed test protocol can unexpectedly affect gait and self-selected walking speed. For example, the treadmill speed controller can induce changes in gait. The mechanics of walking on a treadmill that moves at a constant speed are identical to overground walking. However, when the treadmill accelerates, the belt reference frame is no longer equivalent to a fixed-ground reference [27]. In fact, some belt speed control dynamics can lead subjects to walk at speeds far from their preferred over-ground speed [28]. People may also choose different speeds for different walking tasks, such as to walk for a preset time or a preset distance. If people wish to minimize their energy cost in the fixed distance task, they should walk at a speed close to their normal overground speed. In order to minimize effort in the fixed time task, however, they should walk very slowly or even stand still [3]. Then again, people might not be familiar with the implications of a fixed-time walking task, or might place higher weights on comfort or appearance, or might use a heuristic that defaults to a typical speed in both tasks. The specifics of the task, such as the target distance, may also affect walking speed [29, 30]. People may also change their walking speed in response to other contextual variations, such as the visual environment [31, 32] or auditory cues [33]. Even the details of the verbal instructions provided to participants can have a strong effect on walking speed [34]. Therefore, it is important to validate the self-selected speed test protocol of interest.

A straightforward way of validating a self-selected walking speed test is to compare its measured speeds to those from the standard walking speed test. However, only a few studies have thoroughly compared walking speed on a self-paced treadmill to that during overground walking, and most of those studies were for a motion capture based commercial self-paced treadmill [35, 18]. Van der Krogt and colleagues [35] compared self-selected speeds of typically developing children and children with cerebral palsy in outdoor walking, overground walking in a lab, and walking on a self-paced treadmill in a virtual environment. Children were instructed to “walk at their own preferred, comfortable walking speed.” Both groups of children walked the fastest outdoor, about 5% slower in the lab, and about 10% slower on the self-paced treadmill. Similarly, Plotnik and colleagues [18] compared self-selected speeds in healthy adults during walking for 96 m overground, on a self-paced treadmill, and on a self-paced treadmill with a virtual environment. Subjects were instructed to “walk at their own self-selected preferred comfortable speed.” Subjects walked on the self-paced treadmill at speeds comparable to their overground speeds, while they walked slightly faster when a virtual environment was presented. In addition, walking speed converged faster to steady speed with the virtual environment. These tests demonstrate the value of characterizing response to a self-paced treadmill prior to using it to evaluate the effects of other interventions on self-selected walking speed.

Here, we adapt the force-based self-paced treadmill controller proposed by Feasel and colleagues [15] and evaluate two self-selected walking speed tests using it. First, we explain how the proposed self-pacing controller estimates subject speed and position and adjusts the treadmill speed. Then, we evaluate the speed and position estimations of our controller by comparing them with motion capture data. We then validate the use of the self-paced treadmill for measuring self-selected walking speed. We compare self-selected walking speeds measured from five different speed tests: the standard 10-meter overground walk test, a 150-meter overground walk test, a commonly used manual speed selection treadmill test, a 2-minute self-paced treadmill test, and a 150-meter self-paced treadmill test where subjects can see their goal and progress on a monitor. We compare self-selected walking speed in the 10 m and 150 m overground conditions to test whether the standard measure well represents speeds in longer bouts of walking. We validate the self-paced treadmill tests by evaluating how well they correlate with the standard measure and by comparing them to the commonly used treadmill test. The 2-minute and 150-meter self-paced treadmill tests are compared to each other to examine whether it is necessary to explicitly motivate subjects to walk at their typical speeds by setting target distance and showing their progress. Finally, we discuss the implications of our findings and potential extension of our self-paced treadmill for rehabilitation and assistive device studies.

## Methods

### Self-pacing Algorithm

We revised the self-pacing controller for force-instrumented treadmills proposed by Feasel and colleagues [15]. The central idea is to estimate subject walking speed from foot contact positions and to improve the estimations by incorporating force measurements using a Kalman filter. In our implementation, we track both speed and position with a Kalman filter, which is updated every time step. The filter uses noise matrices determined empirically from motion capture data. We provide a complete description of the algorithm and share the code [36] so that it can be easily used by other researchers.

Our self-pacing controller consists of a subject State Estimator and a treadmill Speed Controller (Fig. 1). The State Estimator takes data from two force plates (third-order Butterworth filter; cutoff frequency: 25 Hz) and the treadmill speed as input and estimates the subject’s speed and position every computational time step, Δ*t*. Based on the estimated speed and position, the Speed Controller adjusts the treadmill speed at the beginning of each footstep.

**Figure 1:**
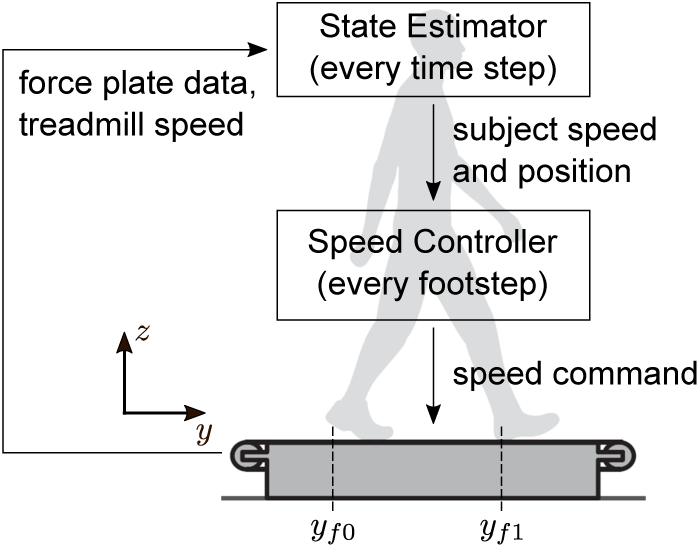
Self-paced treadmill controller. The self-paced treadmill controller consists of a State Estimator and a Speed Controller and only uses force plate data as sensory input.

The State Estimator uses data from the two force plates to measure acceleration, velocity and position of a subject walking on the treadmill and combines the measured values with a Kalman filter. The vertical and fore-aft ground reaction forces (GRFs), *f*_*z*_ and *f*_*y*_, as well as the center of pressure (COP) are calculated from the force-plate data. Foot contact is detected when the vertical GRF exceeds a certain threshold, *f*_*z*_ *> f*_*z*0_ = 20% of body weight. We defined fore-aft foot position on a given step, *y*_*f*1_, as the COP at contact detection. Foot position on the prior step in the lab reference frame, *y*_*f*0_, is calculated by the COP at the previous contact plus the integral of the treadmill speed over the time between the contacts (*y*_*f*1_ and *y*_*f*0_ are shown in Fig. 1). We then estimate the fore-aft acceleration, velocity and position of the subject in the lab reference frame as

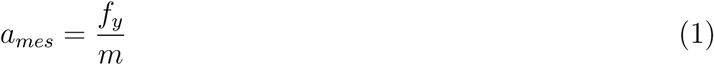

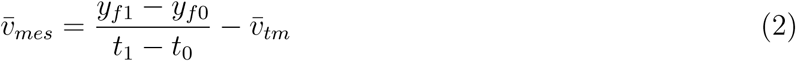

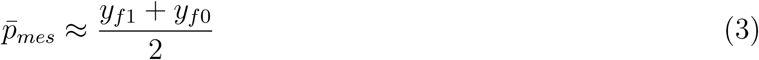

where *m* is the mass of the human subject, and *t*_0_ and *t*_1_ are times when each foot contact occurs, and the variables with a bar indicate mean values during that step (i.e. between consecutive foot contact detections). Eq. 1 is Newton’s second law. Eq. 2 estimates the subject’s mean speed in the lab reference frame, 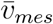, by subtracting treadmill speed 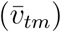 from the subject’s walking speed. The subject’s walking speed is calculated as step length (*y*_*f*1_ − *y*_*f*0_) divided by step time (*t*_1_ − *t*_0_). Eq. 3 defines the subject’s mean position, 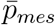, as the middle of the leading and trailing foot placements at a new foot contact.

We implemented a Kalman filter to combine the measurement values *a*_*mes*_, 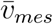 and 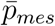 to continuously estimate the subject’s speed and position (Table 1). The filter keeps track of subject speed and position by predicting them every time step from *a*_*mes*_ (Table 1: line 2), and by correcting them with new measurements 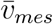 and 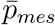 every footstep (line 6). The measurement update is conducted when a new foot contact is detected (line 4). The filter rejects steps of unreasonable duration (greater than 1.2 seconds) to skip the measurement update when subjects cross over the belts (e.g. stepping on the left belt with the right foot). The system model, *A* and *B* (and the observation model *C* = *I*), describes the relationship between the measurement values according to Newton’s second law. The noise matrices, *Q* and *R*, as well as the initial error covariance matrix *P*_0_ are determined from data collected in walking sessions, where two subjects walked on a treadmill at speeds between 0.8 and 1.8 m s^−1^ in ten one-minute trials. The noise matrices are set based on *σ*_*a*_, *σ*_*v*_ and *σ*_*p*_ (Table 1), which are the differences in *a*_*mes*_, 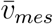 and 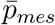, respectively, calculated from force-plate data and motion capture data. *P*_0_ is set to the mean of the values *P* converged to at the end of the pilot sessions.

**Table 1:** Pseudo code of Kalman filter for walking speed and position estimation.

| Pseudo code | Note |
| --- | --- |
| 0: $p_{KF} = 0, v_{KF} = 0, P = P_0$ | initialize |
| 1: **loop** | (every time step) |
| 2: $\begin{bmatrix} p_{KF} \\ v_{KF} \end{bmatrix} = A \begin{bmatrix} p_{KF} \\ v_{KF} \end{bmatrix} + B [a_{mes}]$ | time update:<br>predict speed and position |
| 3: $P = A \cdot P \cdot A^T + B \cdot Q \cdot B^T$ | update error covariance matrix |
| 4: **if** a new foot contact is detected | (every footstep) |
| 5: $K = P \cdot (P + R)^{-1}$ | update Kalman gain |
| 6: $\begin{bmatrix} p_{KF} \\ v_{KF} \end{bmatrix} = \begin{bmatrix} p_{KF} \\ v_{KF} \end{bmatrix} + K \left( \begin{bmatrix} \bar{p}_{mes} \\ \bar{v}_{mes} \end{bmatrix} - \begin{bmatrix} \bar{p}_{KF} \\ \bar{v}_{KF} \end{bmatrix} \right)$ | measurement update:<br>correct speed and position |
| 7: $P = (I - K) \cdot P$ | update error covariance matrix |
| 8: **end if** |  |
| 9: **end loop** |  |

Note that we omitted the observation matrix in lines 5~7 as it is the identity matrix ( $C = I$ ).

The Speed Controller adjusts the treadmill speed to match subject speed and to keep the subject near a baseline position. It updates the treadmill speed once per footstep when a new foot contact is detected. This is different from other self-paced treadmills in previous studies, where speed adjustment is done at a much faster rate (30∼120 Hz) [17, 16, 18]. Controlling the treadmill speed at a higher frequency can lead to undesired dynamics due to natural speed oscillations during walking. Instead of filtering out these oscillations as in the previous studies, we update it at every footstep. Target treadmill speed is set as

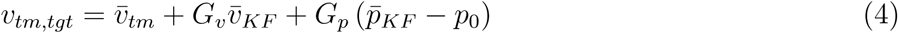

where *p*_0_ is the baseline position, and 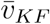 and 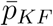 are the subject’s mean speed and position during the last step in the lab reference frame estimated from the Kalman filter. Note that, despite the plus signs, Eq. 4 is a stabilizing negative feedback as the treadmill speeds, *v*_*tm,tgt*_ and 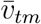, are determined in the opposite direction from the subject speed and position, 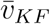 and 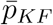, in the lab reference frame. The baseline position *p*_0_ can be predetermined by the experimenter (e.g. *p*_0_ = 0), manually tuned based on subject feedback, or set based on subject data from familiarization trials. In this study, we used the last approach, where we set *p*_0_ for each subject as the average subject position measured during the fixed-speed portion of the treadmill familiarization. In theory, *v*_*tm,tgt*_ with *G*_*v*_ = 1 will be a speed that matches the subject’s estimated walking speed, and *G*_*p*_ = 1 will result in a speed that brings the subject to *p*_0_ in 1 second. However, in pilot tests, we found a controller with these high gains to be unstable. Therefore, we use lower gains of *G*_*v*_ = 0.25 and *G*_*p*_ = 0.1, which we found to be reliable and responsive enough for our study. The treadmill acceleration is set to achieve a target velocity in a certain time as

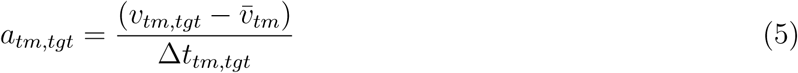

where we use Δ*t*_*tm,tgt*_ = 0.5 sec, similar to the duration of a walking step.

The code of our self-pacing controller and a graphical user interface are publicly available [36]. The self-pacing controller is implemented in Matlab/Simulink Real-Time and runs on a real-time target machine (Speedgoat) at 1000 Hz (i.e. Δ*t* = 0.001). The real-time target machine receives force-plate data from the instrumented treadmill (Bertec) at the same rate. The graphical user interface implemented in Matlab runs on a desktop machine at 100 Hz and allows the experimenter to communicate with the real-time target machine. In addition, it receives the target treadmill speed and acceleration from the real-time target machine and commands it to the treadmill.

### Experiment 1: State Estimator

To evaluate the State Estimator, we compared the estimated position and velocity to those from motion capture data. One subject wore a waist belt with four reflective markers and walked on the force-instrumented treadmill for six one-minute trials. Treadmill speed was manually controlled in most of these trials as we wanted to evaluate the State Estimator independently from the Speed Controller. In the first three trials, the treadmill speed was set to 1.3, 0.8 and 1.8 m s^−1^. In the fourth trial, the treadmill speed changed every 10 sec from 0.8, 1.0, 1.2, 1.4, 1.6 to 1.8 m s^−1^. In the fifth trial, the same speeds were presented in reverse order. Then, the treadmill was controlled with our self-pacing controller in the last trial. Positions of the four reflective markers were captured with a motion capture system (Vicon Vantage; 8 cameras), sampled in 100 Hz and low-pass filtered using a third-order Butterworth filter with a cutoff frequency of 20 Hz. The mean of those maker positions, *p*_*mocap*_, and its time derivative, *v*_*mocap*_, were used for evaluation.

We report how the main outputs of the State Estimator 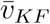 and 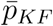 compare to those from motion capture data. For the mean step velocity, we report the root-mean-square (RMS) differences, 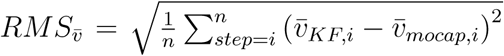, where *n* is the total number of steps in a walking trial, and 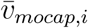 is the mean value of *v*_*mocap*_ on the *i*^*th*^ step. 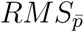 was calculated similarly, but with offset-corrected values for each one-minute trial. This is because 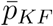 is not tracking the position of the waist. Our approach does not estimate the absolute position of the person’s center of mass, but rather its position relative to the average center of pressure at consecutive foot strikes. Note that any measure of body position can be used to maintain a desirable position on the treadmill by comparing it to a corresponding nominal value, typically determined during a fixed speed calibration trial. In this sense, it is unlikely that any aspect of body position is more useful than any other for self-pacing purposes; only the displacement relative to the nominal position matters.

### Experiment 2: Self-selected Walking Speed Tests

We conducted an experiment to evaluate the validity of our self-paced treadmill in measuring self-selected walking speeds. Ten healthy adults (5 females and 5 males; height: 1.69 ± 0.08 m; age: 25 ± 3 years) participated in the experiment. All subjects participated in a session that consists of familiarization trials and three blocks of five walking speed tests (Fig. 2-a). The familiarization trials were for the subjects to get familiar with walking on our self-paced treadmill and at their comfortable speed in different settings. In addition, the subject’s baseline position, *p*_0_, was found in the fixed-speed portion of the treadmill familiarization. The five walking speed tests in each of the three blocks were presented in random order.

**Figure 2:**
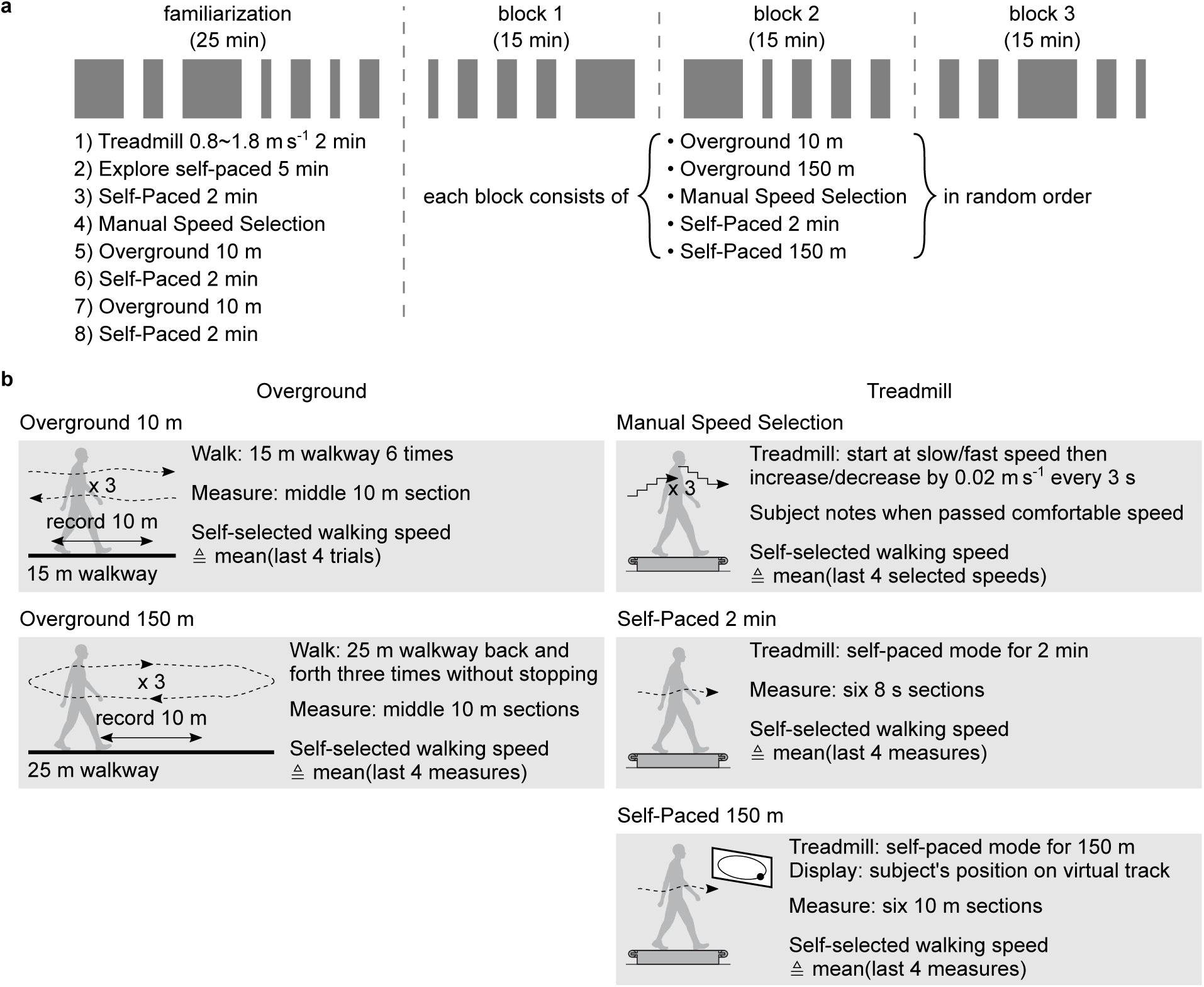
Experimental protocol for self-selected walking speed tests. **a**. The protocol consists of a familiarization session and the main session organized into three blocks. The familiarization session consists of eight overground and treadmill walking trials, which in total takes about 25 minutes. In self-paced treadmill trials, the treadmill first starts at a slow speed, 0.8 m s^−1^, then switches to self-paced mode. Each of the blocks in the main session takes about 15 minutes and consists of five self-selected walking speed tests in random order. **b**. The five walking speed tests consist of two overground and three treadmill tests. In the overground tests, subjects start to walk from standing at the experimenter’s verbal sign “3, 2, 1, go,” and the experimenter measures with a stopwatch the time it takes for the subject to traverse the middle 10 m sections. In the treadmill tests, the treadmill starts at 0.8 m s^−1^ then switches to either speed sweep mode in Manual Speed Selection or self-paced mode in Self-Paced 2 min and Self-Paced 150 m. In Self-Paced 150 m, a monitor shows a 150 m virtual track and a black circle tracking the subject’s position.

We compared five different self-selected walking speed tests. The settings and measurements of the tests are described in Fig. 2-b. Overground 10 m is the standard 10-meter walk test [9, 37] that we used as a reference point in evaluating the outcomes of the other tests. Overground 150 m is to check whether the standard test represents longer distance walking, as walking distance can affect self-selected speed [30]. Manual Speed Selection is a common way to measure preferred walking speed on a treadmill [10, 11, 12]. The correlation between the speed measures in Manual Speed Selection and those in Overground 10 m will be the benchmark value for our self-paced treadmill tests. Self-Paced 2 min and Self-Paced 150 m are the tests using our self-paced treadmill. Subjects were informed whether they would walk for 2 min or 150 m, and, for the latter, subject position was shown on a 150 m virtual track on a monitor in real-time. We applied both fixed-time and fixed-distance tests on the self-paced treadmill to determine whether it was necessary to motivate participants to walk a given distance in order to obtain self-selected walking speeds that correlated well with overground, fixed-distance tasks.

The self-selected walking speed tests were designed to be coherent and comparable with each other. For example, 150 m of walking distance in Self-Paced 150 m was selected to match the distance in Overground 150 m, and the 2 min of walking time in Self-Paced 2 min is the time it takes to walk 150 m at a typical walking speed of 1.25 m s^−1^. Similarly, in Self-Paced 2 min and Self-Paced 150 m, walking speeds were measured in six sections that correspond to the 10-meter-sections in Overground 150 m. We used consistent instructions in all the walking trials [34]. Subjects were instructed to “walk at a comfortable speed” in the overground and self-paced treadmill tests and to verbally indicate when the treadmill gets “faster (or slower) than what you would choose as a comfortable speed” in Manual Speed Selection. When subjects asked for clarification, we elaborated a comfortable speed as “whatever speed feels natural to you.”

We compared self-selected walking speeds measured in each test to the value in the standard over-ground test. The main evaluation was how well walking speed in each test correlated with the speed in the standard test, Overground 10 m. We also compared self-selected speeds in Self-Paced 2 min and Self-Paced 150 m to see whether setting a target walking distance was necessary. In total, we measured 5 sets of 30 self-selected walking speeds: in the five tests, ten subjects walked for three times. For each walking speed test other than Overground 10 m, we report a linear model, *b*_1_ *v*_*OG*10_ + *b*_0_, that fits these 30 measurements to those in Overground 10 m with the minimum mean-squared-error. A test that has a fit of *b*_1_ = 1 and *b*_0_ = 0 indicates that subjects, on average, are likely to walk at the same speed they walked at in Overground 10 m. We also calculate the Pearson’s linear correlation coefficient, *R*, in these pairs of 30 measurements. The correlation coefficient of 1 and 0 correspond to perfect and no correlation, respectively, where a high correlation indicates that much of the variation in measured speeds are captured in the fitted linear model. We considered the linear fit and correlation values to be statistically significant if their *p*-value is smaller than 0.05.

We calculated the variability of self-selected walking speed in each test to determine whether the self-paced treadmill tests were as consistent as the standard overground test. To this end, we calculated the standard deviation of the three walking speed measurements of the same subject within each test, *SD*_*intra*_. We compared these standard deviation values in each test to determine whether certain tests show higher variability than others.

We estimated the time taken to conduct one trial of each walking test to determine whether the self-paced treadmill tests required less time than the common treadmill test. We calculated the minimum time used in all trials in our experiments from the recorded data and report their mean and standard deviation for each walking test. The time for an Overground 10 m trial is calculated as *T*_*OG*10_ = 1.5 × *T*_*OG*10,*rec*_ + 6 × 3, where *T*_*OG*10,*rec*_ is the sum of six recorded times for crossing the 10 m section, multiplication of 1.5 accounts for the additional 5 m walk of the 15 m walkway, and the last term is the three-second countdowns before each of the six bouts. For *T*_*OG*150_ of the Overground 150 m test, we report the recorded time taken by subjects in completing the 150 m course plus 3 s for the countdown. The time used in the Manual Speed Selection, *T*_*MSS*_ is reported as the duration the treadmill was controlled in speed sweep mode plus 3 s for the countdown. Similarly, the times used in Self-Paced 2 min, *T*_*SP*2_, and Self-Paced 150 m, *T*_*SP*150_, are reported as the duration the treadmill was in self-paced mode plus 3 s. Most of the reported times underestimate the actual time required for trials; for example, there were a few additional seconds between each of the six bouts in an Overground 10 m trial, and a few seconds spent before and after speed sweep and self-paced modes in the treadmill trials.

We calculated the time required for walking speed to converge in self-paced treadmill tests to determine the minimum duration of a test with reliable measurements. We observed that participants seemed to converge to steady speed in much less time than the approximately two minutes provided in self-paced walking speed tests. To determine the convergence time in Self-Paced 2 min, we first calculated the mean and standard deviation of walking speeds during the last 20%, or the last 24 sec, of the trial. Then we found the moment when walking speed first entered the range of the mean plus or minus one standard deviation, and determined it to be the convergence time, *t*_*cnvg*_. We determined the convergence distance in Self-Paced 150 m similarly by setting the threshold from the mean and standard deviation of the last 30 m of the trial. Note that the initial treadmill speed was 0.8 m s^−1^ in all the self-paced treadmill trials.

We assessed subject experience in each walking speed test with a survey in order to determine whether the self-paced tests were comfortable and intuitive compared to the standard tests. Subjects rated two written statements for each test after completing all the walking trials. The statements were “it was comfortable walking” and “it was easy to choose my walking speed,”and the subjects had five options: *strongly disagree, disagree, neutral, agree*, and *strongly agree*. We quantified the selections by assigning scores from 1 to 5 for *strongly disagree* to *strongly agree*, respectively.

The statistical significance of differences across walking speed tests, in terms of intra-subject variation, time to measure, and survey scores, was tested using two-way analysis of variance (ANOVA) accounting for different tests and subjects. If a significant effect of test type was found in ANOVA, we conducted paired-sample *t*-test for every pair of tests. We used significance level of *α* = 0.05.

## Results

### Self-pacing Algorithm

The proposed self-pacing controller successfully matched subject speed and kept subjects near the baseline position. In the exploration trial of the familiarization session, all subjects easily walked (or even ran) on the self-paced treadmill at a wide range of speeds (about 0 to 2 m s^−1^).

### Experiment 1: State Estimator

The State Estimator and motion capture system were in close agreement as to the subject speed and position. The RMS differences between estimations of the Kalman filter and motion capture system during the six one-minute trials were 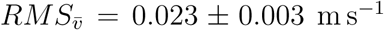 and 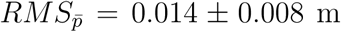. Fig. 3 shows the Kalman filter estimations of the subject speed and position, *v*_*KF*_ and *p*_*KF*_, and their mean values during each step, 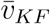 and 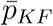, as well as those values from the motion capture data. In addition, the speed and position calculated by merely integrating ground reaction forces are shown to diverge, demonstrating the necessity of the once-per-footstep measurement update of the Kalman filter. Time update using subject acceleration (Table 1: line 2) allows continuous and more accurate tracking of subject speed and position.

**Figure 3:**
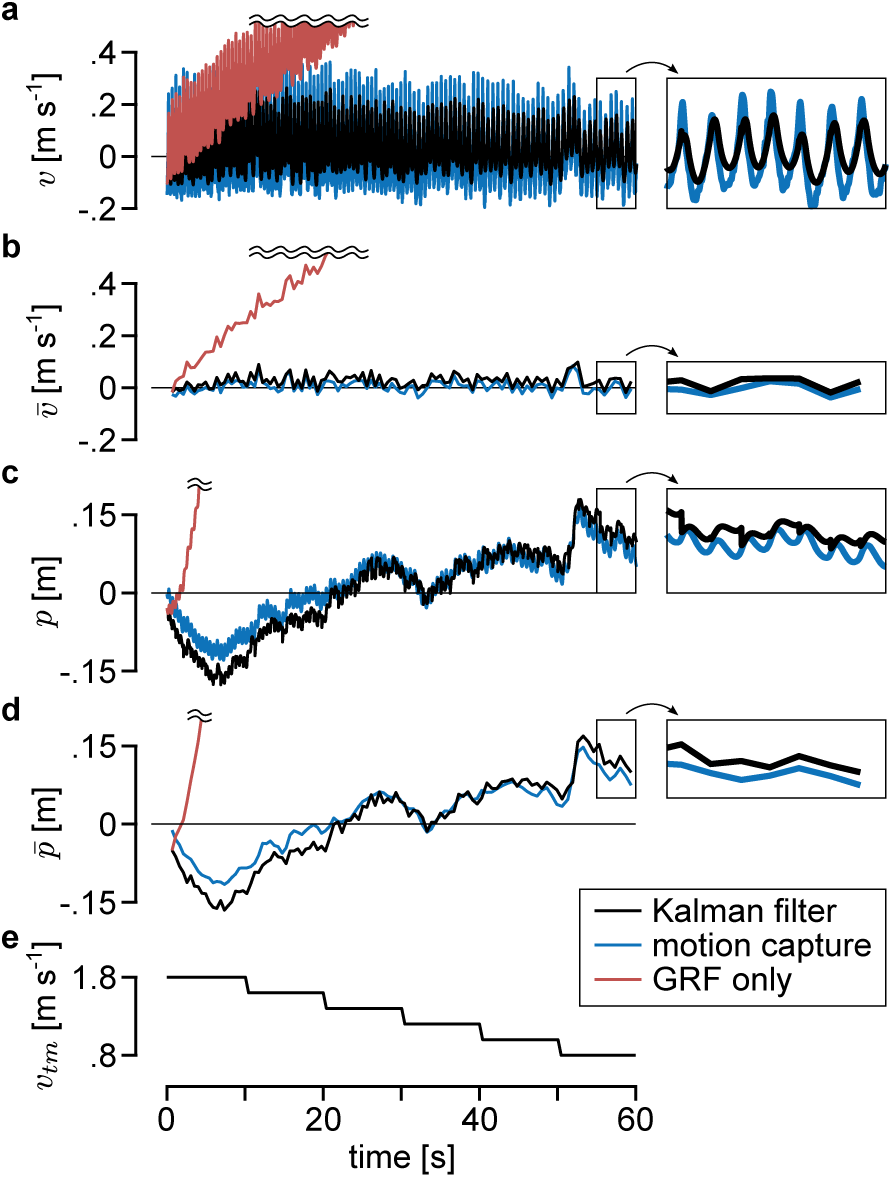
Estimations of State Estimator and motion capture system. The plots show the subject’s estimated **a**. instantaneous speed *v*, **b**. mean speed of each step, 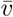, **c**. instantaneous position, *p*, and **d**. mean position of each step, 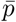. All speeds and positions are in the lab reference frame. The values are estimated with the proposed Kalman filter (black line), motion capture system (blue line), and by simply integrating the ground reaction forces (red line). **e**. The data are collected during a one-minute trial where the treadmill speed, *v*_*tm*_, changes from 1.8 to 0.8 m s^−1^ as shown in the bottom plot.

### Experiment 2: Self-selected Walking Speed Tests

All ten subjects completed the self-selected walking speed test protocol. In the standard Overground 10 m test, the mean and standard deviation of the self-selected walking speeds were 1.32 ± 0.18 m s^−1^, ranging from 0.98 to 1.79 m s^−1^. Leg length, defined as the distance between anterior iliac spine and the medial malleolus, explained 20% of the variance in self-selected walking speed (*R*^2^ = 0.20, *p* = 0.01), which agrees with previous studies [1].

Walking speeds measured in Overground 150 m were close to those in Overground 10 m. The fitted linear model was close to the identity line with a high correlation coefficient (Fig. 4-a). The mean and standard deviation of walking speeds were 1.35 ± 0.19 m s^−1^. This result supports that the standard test, Overground 10 m, reliably measures walking speed in longer distance walking.

**Figure 4:**
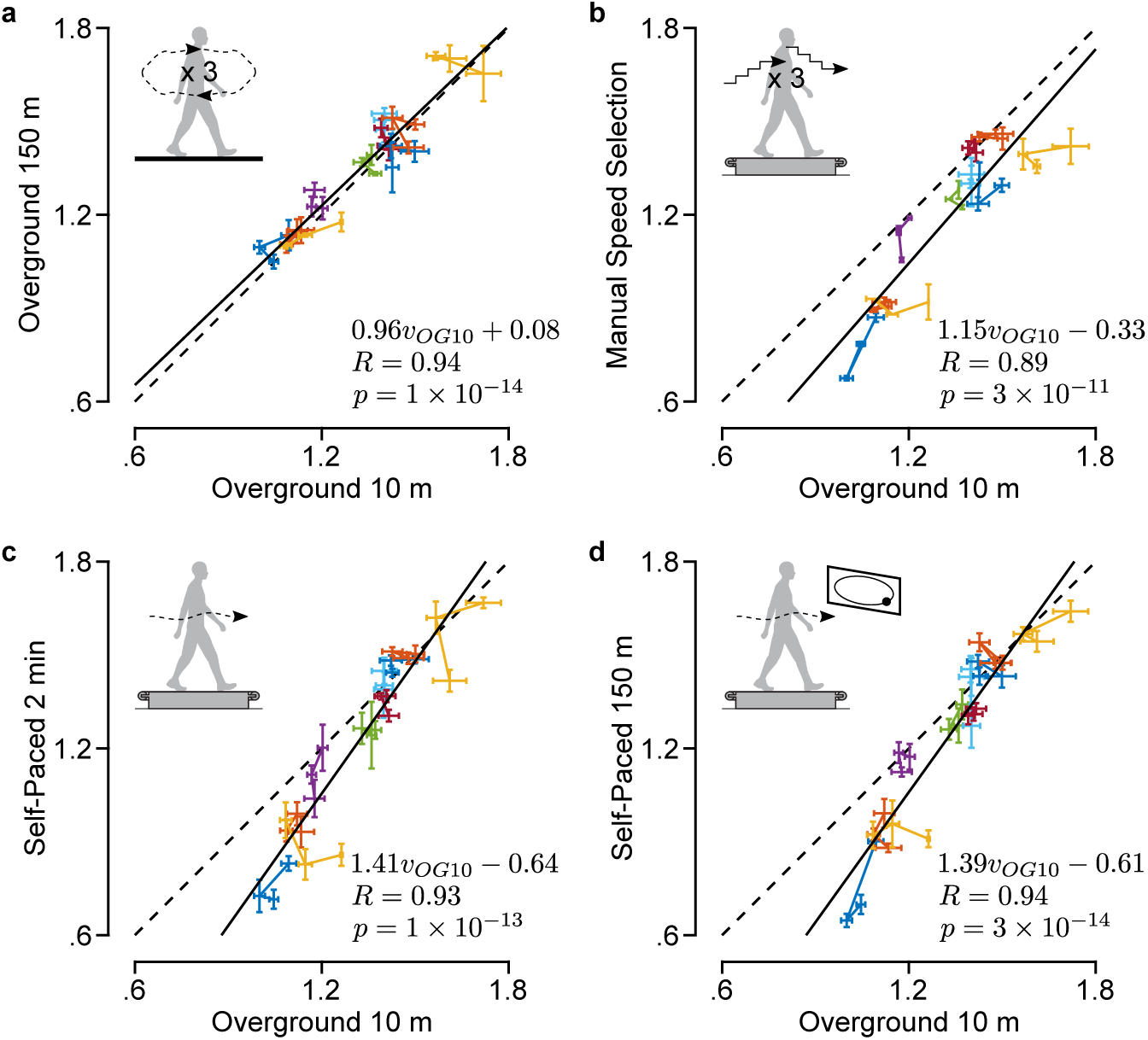
Speeds measured in the self-selected walking speed tests. The self-selected walking speeds measured in **a**. Overground 150 m, **b**. Manual Speed Selection, **c**. Self-Paced 2 min, and **d**. Self-Paced 150 m are compared to those from Overground 10 m. The data points relate a self-selected walking speed measured in a test to the one measured in the standard test in the same block. Each data point is a mean of four measurements (Fig. 2), with whiskers depicting ±1 standard deviation. The exception is for Manual Speed Selection, where the standard deviation is for two measurements because a pair of faster and slower than comfortable speeds are required to obtain one measurement of comfortable speed. Three data points from the same subject are connected with a line and marked in the same color. The linear model, correlation coefficient, and *p*-value for the fit are shown at the bottom right of each plot.

Speeds in Manual Speed Selection were highly correlated with those in Overground 10 m but were slower overall. Walking speeds in Manual Speed Selection were 1.18 ± 0.24 m s^−1^, which was significantly lower (*p* = 0.01) than those in Overground 10 m (Fig. 4-b). This result agrees with previous studies with similar treadmill speed selection tests [10, 12]. The correlation value of *R* = 0.89 between Manual Speed Selection and Overground 10 m is set as the benchmark for our self-paced treadmill tests.

Both Self-Paced 2 min and Self-Paced 150 m were highly correlated with Overground 10 m. The correlation coefficients of the self-paced treadmill tests (*R* = 0.93 and *R* = 0.94) were slightly higher than for Manual Speed Selection (Fig. 4-c and d vs. b). The walking speeds in self-paced treadmill tests were 1.23 ± 0.28 m s^−1^ and 1.23 ± 0.27 m s^−1^, respectively. The speeds were not significantly different from Overground 10 m speeds (*p* = 0.13 in both tests) and were slightly closer than Manual Speed Selection speeds were. However, participants with slower overground walking speeds reduced their speed more on the treadmill. The three slowest subjects walked significantly slower in the self-paced treadmill tests compared to the standard test (0.87 ± 0.11 vs. 1.11 ± 0.07, *p* = 6 × 10^−5^), while the remaining seven subjects did not (1.38 ± 0.15 vs. 1.41 ± 0.13, *p* = 0.49).

Walking speeds measured in Self-Paced 2 min and Self-Paced 150 m were very similar. The fitted model was close to the identity line (*v*_*SP*150_ = 0.96*v*_*SP*2_ + 0.06), and the correlation coefficient was very high (*R* = 0.98, *p* = 7 × 10^−20^).

The intra-subject variabilities in all tests were low and were not significantly different (*p* = 0.49). The average across all tests and participants was *SD*_*intra*_ = 0.042 ± 0.030 m s^−1^. The variability values of individual tests were all lower than 0.1 m s^−1^, which has been suggested as a threshold for clinical significance of differences in walking speed [5, 6, 9].

The self-paced treadmill tests required about a third of the time required for Manual Speed Selection. The mean and standard deviation of the times required for a trial of each test were *T*_*OG*10_ = 87 ± 9 s, *T*_*OG*150_ = 124 ± 16 s, *T*_*MSS*_ = 371 ± 141 s, *T*_*SP*2_ = 125 ± 1 s, and *T*_*SP*150_ = 138 ± 35 s. Walking speed test type had a significant effect on measurement time (ANOVA, *p* = 4 × 10^−37^). All the tests were significantly different from each other (paired *t*-tests, *p* < 0.002), except for Self-Paced 2 min and Self-Paced 150 m (*p* = 0.051) and for Overground 150 m and Self-Paced 2 (*p* = 0.754). Manual Speed Selection took the longest on average and also was the most variable across subjects. The large time variation was due to some subjects having large gaps between the speeds identified to be faster or slower than comfortable speeds while others had smaller gaps.

Analysis of speed convergence in the self-paced treadmill tests suggests that the preset time and distance can be much shorter than 2 min and 150 m. The mean and standard deviation of the convergence time in Self-Paced 2 min were *t*_*cnvg*_ = 22 ± 22 s while mean and standard deviation of convergence distance in Self-Paced 150 m were *d*_*cnvg*_ = 42 ± 29 m (Fig. 5). The convergence distance in Self-Paced 150 m, *d*_*cnvg*_, corresponded to *t*_*cnvg*_ = 34 ± 22 s in time, significantly longer than in Self-Paced 2 min (*p* = 0.048). This result suggests that the times used in the current Self-Paced 2 min (*T*_*SP* 2_ = 125 s) and Self-Paced 150 m (*T*_*SP*150_ = 138 s) could be much shorter. For example, the average speed during the last five seconds of the first minute of the Self-Paced 2 min test is not statistically different from the current measure (*p* = 0.89). This would require about one sixth the time of the conventional treadmill speed test.

**Figure 5:**
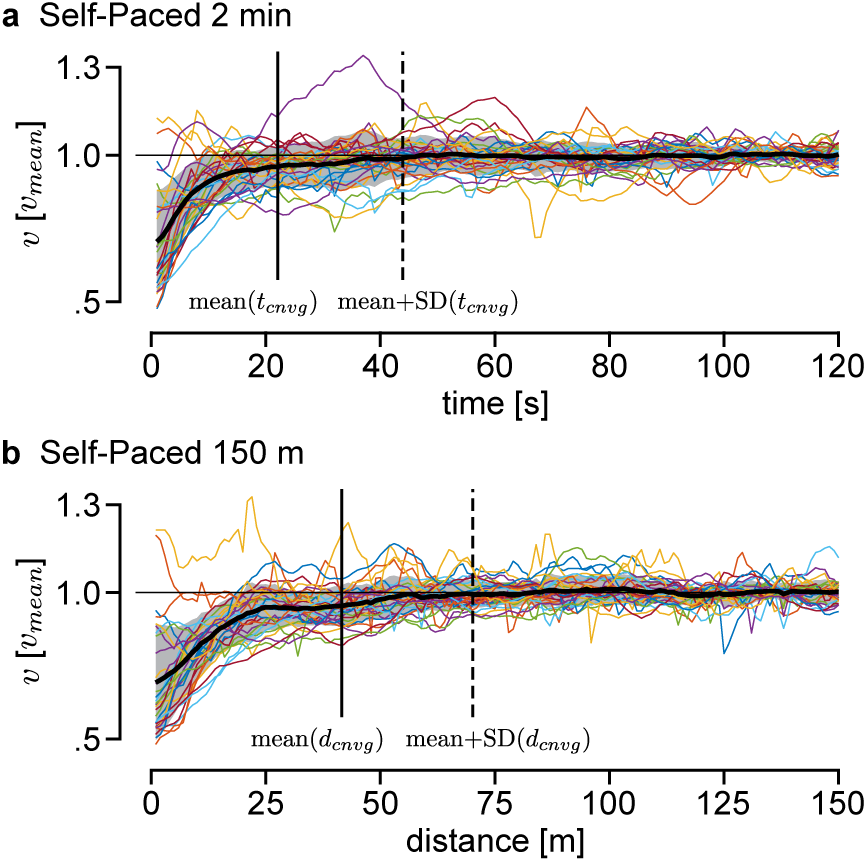
Convergence of walking speeds in self-paced treadmill tests. Walking speeds normalized by final estimated speed in **a**. Self-Paced 2 min and **b**. Self-Paced 150 m tests. Walking speed from individual trials are shown in colored lines. The mean and ± 1 standard deviation across all trials are shown as a black line and gray shaded area. The solid and dotted vertical lines indicate the mean and mean plus one standard deviation of convergence time and distance.

The survey results suggested that subjects found walking at their comfortable speeds in the self-paced treadmill tests to be as comfortable as in the common treadmill speed selection test but not as comfortable as in overground tests. The mean and standard deviation of the scores for “it was comfortable walking” were 4.3 ± 0.7 for Overground 10 m, 4.4±0.5 for Overground 150 m, 3.5 ± 1.0 for Manual Speed Selection, 3.9 ± 0.7 for Self-Paced 10 m, and 3.8 ± 0.8 for Self-Paced 150 m, where 1 is *strongly disagree* and 5 is *strongly agree*. The scores for the “it was easy to choose my walking speed” statement were 4.4 ± 0.7, 4.5 ± 0.7, 3.0 ± 1.2, 3.3 ± 0.7 and 3.4 ± 1.0, respectively. Speed test type had a significant effect on survey results (ANOVA, *p* = 0.002 and 1 × 10^−5^, respectively). Comfort and ease of speed selection in self-paced tests were not significantly different from those in the conventional treadmill test (paired *t*-tests, *p >* 0.10) but were worse than those in overground tests (*p* < 0.053).

## Discussion

Our results indicate that the proposed self-paced treadmill can be used to measure self-selected walking speed. Subjects selected walking speeds in both self-paced treadmill tests that were highly correlated with their speeds in the standard overground test. Intra-subject speed variations in the self-paced treadmill tests were low, demonstrating repeatability. The self-paced treadmill tests required only about a third of the time to complete of a common treadmill test, with no reduction in comfort or ease.

Although the walking speeds from self-paced treadmill tests highly correlated with the standard 10-meter walk test, the actual speeds were not the same. More specifically, subjects who walked at slow speeds in Overground 10 m walked even slower in Self-Paced 2 min and Self-Paced 150 m (Fig. 4-c,d). We can speculate different reasons for this observation. First, our self-pacing controller may be tuned better for normal and fast walking than walking at slow speeds. However, that would not explain why the slow walking subjects also selected slower speeds in Manual Speed Selection (Fig. 4-b). Second, which is more compelling in our opinion, contextual changes [31, 32, 33] other than segment dynamics (i.e. force interactions between subjects and the treadmill or ground) may have a larger effect during slower walking. The influence of these contextual changes may depend on walking speed because control strategies may change for different speeds [38, 39] as modeling studies suggest slower walking should rely more on active balance control than on passive dynamics [40]. This hypothesis could be tested by studying how the amount of context-induced gait changes correlate with walking speed. Whatever the reason, the strong correlation between self-paced and overground speeds suggests that changes in self-selected walking speed on the self-paced treadmill will translate into changes during overground walking, though the absolute magnitudes may differ.

Subjects selected to walk at very similar speeds on our self-paced treadmill whether they were walking for a preset time or a preset distance. This was unexpected because it would seem inconsistent with the minimum effort principle. So why did subjects walk at similar speeds in the preset time (Self-Paced 2 min) and present distance (Self-Paced 150 m) tests? First, subjects may have tried to fulfill the experimenter’s expectation. We instructed the subjects to walk at their comfortable speed in all five tests, which the subjects may have interpreted as walking at a particular speed. However, such interpretation or intent of matching experimenter expectation was not apparent from subject feedback. Second, it could be that the objective of walking for a preset time was not clear to subjects because it is different enough from other walking tasks that they had experienced. Walking for a preset distance is close to walking to a target location, which is very common in daily life. Walking or running on a treadmill in a gym for a preset time as a workout might seem similar but is different from the preset time test in our study, in that the speed is usually set based on energy expenditure goals. For the unique task of walking for a preset time in an experiment, subjects may have aimed to walk in a way they were most familiar with, which is to walk for a preset distance. Regardless of the reason, all subjects in our study self-selected to walk at similar speeds in the preset time and preset distance tests. Therefore, we can use the preset time on a self-paced treadmill to measure self-selected walking speeds, which can be easier to administer than for preset distance.

The proposed self-pacing controller is different from most previous controllers in that it uses data from treadmill force plates to estimate subject speed and position. Therefore, it requires a force-instrumented treadmill, and subjects should not cross over the belts when stepping, which can interfere with their natural gait. However, stepping on the correct belt on an instrumented treadmill is a common requirement for gait studies [13], in which case, the self-pacing controller can be used with little overhead. We have previously tested other approaches that require additional parts on subjects, such as motion capture markers or string potentiometers, and those setups can easily increase the burden in complex gait experiments, such as studies on robotic exoskeletons or prostheses [14, 41]. A useful future extension in this direction is improving the self-pacing controller to work with a single force-plate, which would allow subjects to cross over the belts.

Another difference from most prior self-pacing controllers is that ours adjusts the treadmill speed only once per footstep. Most other self-paced treadmill controllers update treadmill speed at a higher frequency (30∼120 Hz) [17, 16, 18]. If the treadmill speed instantaneously matches subject body speed, it will fluctuate within every stride due to natural speed oscillations in normal walking (Fig. 3-a) and may introduce undesired treadmill dynamics. To minimize this effect, previous studies low-pass filtered the estimated body state with a low cutoff frequency (e.g. 2 Hz), which can introduce time delays. Instead, our controller updates the treadmill speed once-per-footstep based on the mean values in that footstep. We find our approach to be conceptually more consistent with the control goal of matching walking speed, not instantaneous speed. A more thorough investigation of treadmill speed adjustment strategies could be instructive and might improve the self-pacing controller. For example, we use a simple heuristic control scheme (Eq. 4) with low control gains in matching subject speed and position, which is similar to previous approaches [16]. While higher gains can respond more quickly to speed and position changes, we empirically found lower gains to be stable and reliable for walking at steady speeds and moderate speed changes. Gain scheduling that matches large speed changes as well as steady walking would extend the potential use of self-paced treadmills in gait studies.

The proposed self-paced treadmill can be used in rehabilitation treatment and in gait assistance research but should be re-validated for substantially different populations or tasks. All of the subjects that participated in our experiment found walking on the self-paced treadmill intuitive and easy. However, the subtle dynamics and apparent contextual differences induced by self-paced treadmills may have a larger effect for subjects with different health status or for different locomotion tasks. For example, it has been reported that children with cerebral palsy experienced larger changes in gait on a self-paced treadmill than typically developing children [35]. Nevertheless, for healthy adults walking at typical speeds, self-selected walking speed on this self-paced treadmill can be used as an indication of overground walking behavior.

## Conclusions

We presented a self-paced treadmill controller for force-instrumented treadmills that can be used to measure self-selected walking speeds. The controller is adapted from a previous study [15] and solely uses force-plate data to estimate and adapt to the subject’s walking speed and position. To validate its use for measuring self-selected walking speeds, we compared walking speeds measured in a range of walking speed tests, where the subjects were instructed to walk at or select their comfortable speed. The tests using our self-paced treadmill measured walking speeds that were highly correlated with those from the standard overground test. The differences in the measured speeds from the self-paced treadmill and overground tests were small but consistent. The low intra-subject variability of measured speeds supports the reliability of the self-paced treadmill tests. The times required for the self-paced treadmill tests were a few times less than that for a common treadmill test, where subjects manually select their comfortable speeds, with the potential for further substantial reductions in duration. Subjects found the self-paced treadmill tests to be as comfortable and easy as the common treadmill test. These results demonstrate that measurements of self-selected walking speed made using the self-paced treadmill are relevant to overground conditions, and that the self-paced treadmill provides a strong alternative to manual speed selection on an instrumented treadmill. We provide a complete description and code for the self-pacing controller and graphical user interface to facilitate use by other gait researchers and clinicians [36].

## Abbreviations

ANOVA: analysis of variance;
GRF: ground reaction force;
COP: center of pressure;
RMS: root mean square

## Acknowledgements

The authors thank all participants of this study as well as Maxwell Donelan and Arthur Kuo for discussions about preset time and preset distance walking.

## Author’s contributions

SS and SC conceived the study and designed the experiment, SS and HC developed the algorithm, SS conducted experiments and analyzed data, SS drafted the manuscript, SS and SC edited the manuscript, and all authors approved the submitted manuscript.

## Availability of data and materials

All data collected in the study are available from the corresponding author on reasonable request. The code for self-paced treadmill is available in the self-paced-treadmill repository on GitHub [36].

## Ethics approval and consent to participate

Ethical approval for the study was granted by the Stanford University Institutional Review Board. All participants provided written informed consent.

## Consent for publication

Not applicable.

